# Death is on Our Side: Paleontological Data Drastically Modify Phylogenetic Hypotheses

**DOI:** 10.1101/723882

**Authors:** Nicolás Mongiardino Koch, Luke A. Parry

## Abstract

Fossils are the only remaining evidence of the majority of species that have ever existed, providing a direct window into events in evolutionary history that shaped the diversification of life on Earth. Phylogenies underpin our ability to make sense of evolution but are routinely inferred only from data available from living organisms. Although extinct taxa have been shown to add crucial information for inferring macroevolutionary patterns and processes including ancestral states, paleobiogeography and diversification dynamics, the role that fossils play in inferring the tree of life itself is controversial. Since the early years of phylogenetic systematics, different studies have dismissed the impact of fossils due to their incompleteness, championed their ability to overturn phylogenetic hypotheses or concluded that their behavior is indistinguishable from that of extant taxa. Here we show paleontological data has a remarkable effect in phylogenetic inference. Fossils often have higher levels of topological influence than extant taxa, while inducing unique topological rearrangements. Previous studies have proposed a suite of explanations for the topological behavior of fossils, such as their retention of unique morphologies or their ability to break long branches. We develop predictive models that demonstrate that the possession of distinctive character state combinations is the primary predictor of the degree of induced topological change, and that the relative impact of taxa (fossil and extant) can be predicted to some extent before any analysis. Our results bolster the consensus of recent empirical studies by showing the unique role of paleontological data in phylogenetic inference, and provide the first quantitative assessment of its determinants, with broad consequences for the design of taxon sampling in both morphological and total-evidence analyses.

---

The overwhelming majority of species produced through the diversification of life on Earth are now extinct (Simpson 1952; Raup 1992). Even though much of this diversity is lost, a significant proportion has been preserved in the fossil record, which often provides the most direct evidence of evolution in deep-time. Although evolutionary inferences are routinely performed using data from extant taxa alone, this can often provide only a partial, or even misleading, view of evolutionary processes and patterns. Consequently, paleontological data can not only expand the range of evolutionary questions accessible to inquiry, but also drastically improve estimates of evolutionary phenomena. Both simulations and empirical case studies attest to the positive effect that incorporating extinct diversity can have on the reconstruction of ancestral states (Finarelli and Flynn 2006; Finarelli and Goswami 2013; Puttick 2016), rates and modes of macroevolution (Slater et al. 2012; Bokma et al. 2015; Mitchell 2015; Schnitzler et al. 2017), diversification dynamics (Liow et al. 2010; Quental and Marshall 2010; Rabosky 2010; Mitchell et al. 2018) and historical biogeography (Wood et al. 2012; Field and Hsiang 2018). Fossils also provide the most direct and widely employed evidence used to time-calibrate phylogenies (Laurin 2012; Dos Reis et al. 2016), a key first step in most of modern comparative biology.

Despite the key role of the fossil record in understanding evolutionary history, the degree to which extinct taxa contribute to the inference of phylogenetic relationships has been much more controversial. Although this is often discussed in the context of the merits (and caveats) of reconstructing phylogeny using morphology (e.g., Scotland et al. 2003; Jenner 2004), the debate has a longer history. Hennig (1966) first suggested that the higher proportion of missing data in fossils should compromise their usefulness for elucidating phylogenetic relationships, a view shared by other early systematists (Løvtrup 1977; Ax 1987). From this perspective, fossils hold a subsidiary role, and their significance should only be discussed in light of phylogenies built from extant taxa (Patterson 1977; Nelson 1978). This assumes, either implicitly or explicitly, that fossils do not modify tree topology (Hennig 1981; Patterson 1981; Goodman 1989), and can be grafted onto phylogenies inferred using data from living species.

However, extinct organisms have several characteristics that should make them especially important for inferring accurate trees from morphological data. Fossils allow taxon sampling to be extended beyond the reach of molecular data, and preserve character state combinations not present among living clades, potentially modifying homology statements, character polarity and tree rooting (Doyle and Donoghue 1987; Marshall and Schultze 1992; Novacek 1992a; Wilson 1992; Smith 1994, 1998; Forey and Fortey 2001; Edgecombe 2010). Fossil terminals can also occupy unique phylogenetic positions, lying close to divergence events, in the midst of ancient and rapid radiations, or subdividing the long branches that often separate morphologically distant extant lineages (Doyle and Donoghue 1987; Gauthier et al. 1988; Donoghue et al. 1989; Huelsenbeck 1991; Sumrall 1997; Smith 1998; Wills and Fortey 2000; Smith and Turner 2005; Mayr 2006). Their morphology may often resemble that of the common ancestors from which extant clades originated, being less modified by subsequent evolutionary history (Beck and Baillie 2018; Asher et al. 2019). Furthermore, the proportion of missing data does not necessarily compromise the phylogenetic placement of terminals, nor the overall resolution of phylogenetic analyses (Kearney and Clark 2003; Wiens 2003a, b; Prevosti and Chemisquy 2010; Pattinson et al. 2014), and incomplete terminals can in fact increase topological accuracy (Huelsenbeck 1991; Wiens 2005). Fossils are therefore expected to have a strong topological impact, and early claims for their dismissal were rapidly falsified by several case studies (Gauthier et al. 1988; Donoghue et al. 1989; Doyle and Donoghue 1992; Novacek 1992b; Wilson 1992; Cloutier and Ahlberg 1995; Smith 1998).

Nevertheless, it is difficult to draw general conclusions regarding the impact of paleontological data from individual studies, and it further remains unclear whether fossil terminals modify phylogenetic trees above a baseline of expected change given increased taxon sampling. The only empirical study addressing these issues analyzed 45 empirical morphological matrices and concluded that there was no significant difference in the degree of induced topological change between fossil and extant terminals (Cobbett et al. 2007). This conclusion was supported by first-order taxon jackknifing experiments (i.e., comparison of topologies obtained with and without a focal taxon). Although the authors interpreted this result as strongly supporting the inclusion of fossils in phylogenetic analyses, this was mostly justified in the lack of a distinctive behavior by paleontological data. This conclusion not only conflicts with the literature cited above, much of which considers paleontological data to be unique (in either a negative or positive way), but also with a number of more recent studies where prominent and long-standing cases of conflict between morphological and molecular trees have been claimed to be resolved through the addition of key fossils (Legg et al. 2013; Parry et al. 2016; Coiro et al. 2018; Simões et al. 2018; Miyashita et al. 2019). However, two points should be noted regarding this discrepancy: 1) Given their experimental design, Cobbett et al. (2007) never tested the degree to which fossils overturn relationships inferred *exclusively* from living taxa, and 2) recent case studies do not necessarily claim that fossils have a strong topological effect, rather that the type of change induced is not generated by increasing sampling among extant taxa. These two aspects of the interaction between paleontological and neontological data in phylogenetic studies have never been systematically explored.

Given that relationships among living clades are now routinely inferred from molecular data, this discussion has been deemed obsolete (Scotland et al. 2003). However, even in the genomic era, morphology will remain the only means to resolve the relationships among extinct species, as well as their position relative to extant clades (Giribet 2015; Lee and Palci 2015). Incorporating the information preserved in the fossil record into phylogenetic frameworks not only greatly improves the behavior of phylogenetic comparative methods (Slater and Harmon 2013; Goswami et al. 2016; see above), but also allows the use of tip-dating approaches to divergence time estimation, which require fewer assumptions and make better use of stratigraphic data than more traditional node-dating methods (Ronquist et al. 2012a; Heath et al. 2014; Lee and Palci 2015; Zhang et al. 2016). Furthermore, it has been shown that morphological data has the power to modify tree topology in total-evidence analyses, even when constituting a minimal fraction of the data (Wiens et al. 2010; Bapst et al. 2018; Cascini et al. 2019). Moreover, phylogenies inferred from molecular data are far from stable for all nodes in the tree of life, with several recent examples of phylogenomic data generating conflicting topologies that have sparked controversy (e.g., Ballesteros and Sharma 2019; Philippe et al. 2019). Therefore, morphological data remains an important source of independent data, and understanding the impact of including fossil taxa in phylogenetic analyses remains paramount to obtaining a complete and accurate picture of evolutionary history.

Here we employ multiple empirical large-scale morphological matrices to explore the degree and type of topological change exerted by fossils on trees of extant lineages. As probabilistic approaches to morphological inference have become increasingly common, we extend previous efforts by analyzing results obtained under both maximum parsimony (MP) and Bayesian inference (BI). Finally, we evaluate for the first time several potential determinants of the topological impact of taxa, allowing us to build a framework that can predict whether a terminal will have a strong effect on tree topology.

## MATERIALS & METHODS

### Dataset Selection and Subsampling Procedure

Datasets were selected for their size, large number of fossil and extant taxa, and relatively small proportions of missing data. We focused on large-scale empirical morphological matrices (also known as ‘phenomic’ matrices (O’Leary and Kaufman 2011; O’Leary et al. 2013)), as these are expected to generate better constrained distributions of optimal topologies. Datasets were also required to contain reasonably large numbers of both fossils and extant taxa, therefore allowing for the comparison of their topological effects. Given that some of our analyses explored the topological changes induced on trees built *only* from extant terminals, high numbers of these were especially important. Finally, fossil taxa had to be coded for a significant fraction of the total number of characters. We translated these requirements into a set of rules, employing matrices that had: 1) a number of characters larger than the number of taxa; 2) at least 40 extant terminals; 3) at least 20 fossil terminals; 4) a fraction of missing data among fossils less than 80%. Six datasets (shown in Table 1 and available as SI File 1) satisfied these criteria, and were thus used for all analyses. Preliminary analyses with a larger number of datasets revealed that either the subsampling procedure or the subsequent statistical analyses (see below) could not be performed if these requirements were not enforced. In each case, a single outgroup was included to root all trees. Characters considered ordered by the authors were analyzed as such. If datasets were modified in any other way, details can be found in SI File 2.

**Table 1.** Morphological matrices employed differ in taxonomic scope, character coding strategies and size.

| Dataset | Number of<br>ingroup taxa | Percentage<br>of fossils | Number of<br>characters | Percentage of multistate<br>– ordered characters | Percentage of missing<br>data (extant – fossil) |
| --- | --- | --- | --- | --- | --- |
| Cocomorpha (Vea and Grimaldi 2015) | 116 | 37.1 | 174 | 50.6 – 0.0 | 23.8 – 50.2 |
| Echinoidea (Kroh and Smith 2010) | 163 | 55.8 | 303 | 34.0 – 5.3 | 17.5 – 28.0 |
| Lemuriformes (Herrera and Dávalos 2016) | 62 | 32.3 | 421 | 69.1 – 0.0 | 44.0 – 71.1 |
| Mammalia (O’Leary et al. 2013) | 83 | 44.6 | 4541 | 21.2– 0.0 | 39.4 – 65.8 |
| Panarthropoda (Legg et al. 2013) | 309 | 69.2 | 753 | 14.2 – 0.0 | 55.2 – 76.1 |
| Squamata (unpublished, based on (Gauthier et al.<br>2012, Mongiardino Koch and Gauthier 2018)) | 198 | 28.3 | 767 | 35.3 – 23.9 | 32.0 – 62.6 |

Datasets were imported into the R statistical environment (R Core Team 2019) using function ReadMorphNexus from package *Claddis* v. 0.3 (Lloyd 2016). For each dataset, 25 initial pseudoreplicated matrices composed of *n* randomly selected extant taxa were generated (Fig. 1, step 1), and phylogenetic inference was performed before and after the incorporation of further terminals. Unlike previous efforts that measured the topological impact of adding individual terminals (Cobbett et al. 2007), we explored the topological effects induced by the simultaneous addition of groups of terminals (of size *m*) to these initial replicates (Fig. 1, step 2). We believe this approach to more accurately reflect the way in which morphological datasets grow with time, as well as providing greater subsampling flexibility and increased statistical power. The values of *n* and *m* were determined for each matrix following the approach described in SI File 2.

**Figure 1:**
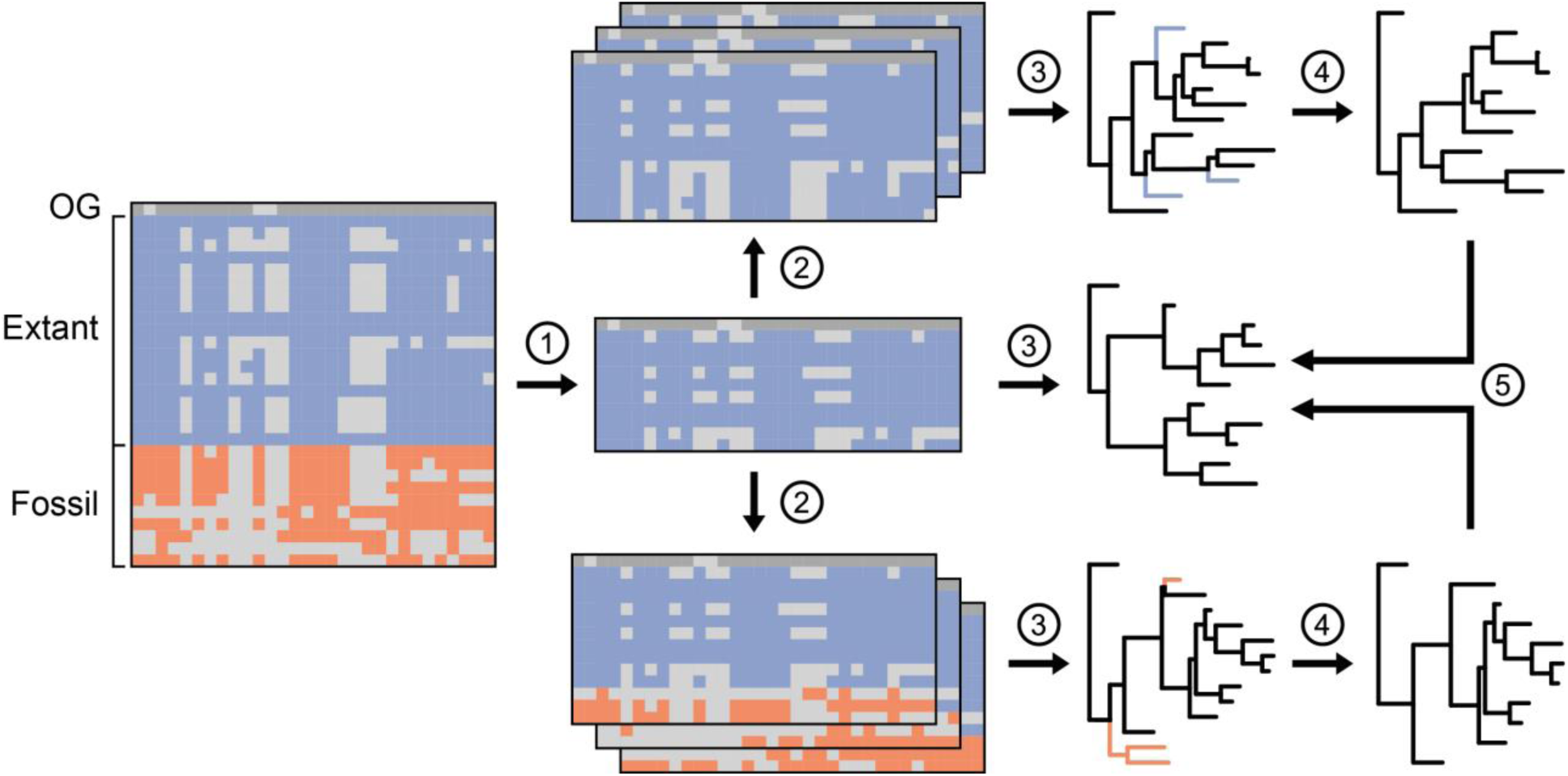
Protocol for the assessment of topological impact. **1.** Matrices were subsampled to a fixed number (*n*) of extant taxa plus one outgroup (OG). 25 such initial replicates were built from each dataset. **2.** A given number (*m*) of terminals were added to these initial replicates. These were composed entirely of extant (top, blue), fossil (bottom, orange), or pseudoextinct taxa (not shown). Three iterations of taxon addition were performed. **3.** Matrices were subject to phylogenetic analysis under maximum parsimony (MP) and Bayesian inference (BI), resulting in a set of optimal topologies from which at most 100 were sampled at random (represented here by a single tree). **4.** Topologies obtained after taxon addition were pruned to match the taxon sampling of the initial replicate. **5.** Topologies were compared using Robinson-Foulds (RF) distances. Further rounds of addition of *m* terminals were performed on the matrices resulting from step 2, followed by the same subsequent steps. The values of *n* and *m* were determined for each dataset as explained in the SI File 2. The same color scheme is used throughout.

The groups of added terminals were of one of three different types: fossil, extant and pseudoextinct, with the last two providing different bases with which to compare the effects induced by fossils. The first two of these groups were generated by selecting at random among the fossil or extant taxa left unsampled. However, the direct comparison of the topological effect of fossil and extant terminals might be confounded by the systematic difference in the amount of data coded between them (see Table 1). Therefore, a third group of terminals (pseudoextinct, following the nomenclature of Springer et al. 2007) was generated by selecting at random *m* of the unsampled extant taxa and pairing each with a randomly selected fossil. Characters missing in the fossil were then deleted from the extant terminal with which it was paired (see Pattinson et al. 2014 for an equivalent approach). This procedure preserves the pattern of missing data found in fossils, and is expected to generate better fossil analogs compared to the deletion of random characters given how morphological structures differ in both their preservation potential and phylogenetic signal (Sansom and Wills 2013; Mounce et al. 2016; Sansom and Wills 2017; Sansom et al. 2017). Note however that the total amount of data in pseudoextinct taxa is expected to be slightly lower than that of fossils, as extant taxa already have missing data that might be coded in some fossils. Comparison of the amounts of missing data between these three groups of terminals can be found in Figure S1 (SI File 3). While fossils have on average 10.5%-30.9% more missing data than extant taxa (depending on the dataset), pseudoextinct taxa have on average 4.6%-8.5% more missing data than fossils. This represents a 2.3-4.6 times reduction in the discrepancy of missing data between compared groups. Given how missing data is known to affect the topological impact of terminals (Huelsenbeck 1991; Wiens 2003b, 2005), pseudoextinct terminals likely provide a better baseline for the topological effect that can be expected from fossils.

For each of the 25 initial replicates per dataset, terminals of the different types were added in a stepwise manner in groups of size *m* until unsampled terminals were exhausted. Furthermore, three iterations of this procedure were performed in order to estimate the average effect of taxon addition to a given taxonomic composition. This entire subsampling procedure generated between 2,275 and 3,625 morphological matrices per dataset.

### Phylogenetic Inference

Phylogenetic inference on these matrices was performed under both MP and BI (Fig. 1, step 3). Although MP was historically favored as a method of inference from morphological data, several studies have argued probabilistic approaches of inference, and more specifically BI under variants of the Mk model (Lewis 2001), might outperform parsimony-based methods (Wright and Hillis 2014; O’Reilly et al. 2016; Puttick et al. 2017). These results have received different criticisms (Goloboff et al. 2018, 2019; Smith 2019), but ultimately rely on analyses performed on simulated data, under conditions where the accuracy of the inference can be evaluated. It still remains unclear whether these results can be extrapolated to the analysis of empirical matrices (Sansom et al. 2018; Schrago et al. 2018; Goloboff et al. 2019), but it can be argued that they have established BI under the Mk model as a valid alternative to MP. Furthermore, given the flexibility of Bayesian methods to combine different data sources and calibrate divergence times, BI of morphological datasets is likely to become common in the future. We here treat these as valid methods of inference from morphological data and explore whether any of our results varies depending on method choice, extending previous attempts which had relied exclusively on MP (Cobbett et al. 2007).

Inference under MP was performed using TNT 1.5 (Goloboff and Catalano 2016) under equal weights, using driven tree searches with five initial replicates that were subject to new technology search heuristics (Goloboff 1999; Nixon 1999). Search was continued until minimum length was found twenty times. TBR branch swapping was then performed on the topologies in memory, retaining a maximum of 10,000 maximum parsimony trees (an example TNT batch script to perform tree searches can be found as SI File 4). Bayesian analyses were performed in MrBayes 3.2.6 (Ronquist et al. 2012b) under the Mk+Γ model with a correction for coding only parsimony-informative characters (MK_parsinf_). Two runs of four Metropolis-coupled MCMC chains were continued for either one million generations or until a standard deviation of split frequencies < 0.01 was attained. We considered this condition to represent an accurate sampling of the posterior distribution of topologies (as have other authors; e.g., Puttick et al. 2019). Trees were sampled every 100 generations and the initial 50% was discarded as burn-in, therefore retaining a maximum of 10,000 posterior topologies.

### Estimating and Analyzing Topological Impact

Optimal topologies from MP and BI runs were imported into R using functions from packages *ape* (Paradis and Schliep 2018) and *TNTR* (Matzke 2015), and a random sample of up to 100 topologies was drawn from each analysis. Topologies before and after the addition of groups of taxa were compared directly by calculating their normalized Robinson-Foulds (RF) distance (Robinson and Foulds 1981) after pruning to the same taxonomic composition (Fig. 1, step 4 and 5). All trees analyzed were thus fully bifurcating, avoiding the issue of computing similarity when topologies differ in their level of resolution (Smith 2019). For pairs of unrooted bifurcating trees the RF distance is equivalent to the number of bipartitions not shared (Penny and Hendy 1985), and was normalized to the total number of bipartitions in both. RF distances were computed using package *phangorn* v. 2.5.5 (Schliep 2011). In order to derive a single metric that reflects the distance between sets of trees, we measured both the mean and minimum RF distance of a given tree to all trees in the other set, and took the average of the resulting values. Given the strong linear correlation between these two estimates (Fig. S2 of SI File 3; all Pearson’s *r* > 0.89, *P* < 10^−259^), analyses employed the average of distances to the nearest neighbor. This metric was favored as it attains a value of 0 if the sets being compared are identical (Cobbett et al. 2007), and allows for a more straightforward comparison of sets of trees derived from MP and BI, which are expected to systematically differ in their within-set distances (O’Reilly et al. 2016; Schrago et al. 2018).

We explored the degree of topological change generated by taxon addition by summarizing RF distances across iterations and analyzing them with generalized linear models. The effect produced by the incorporation of fossils to topologies of extant taxa was compared in multiple ways with that introduced by further increasing sampling of extant taxa. First, we used the number (log-transformed) and type (fossil/extant/pseudoextinct) of added terminals as predictors of topological change. Extant and pseudoextinct terminals provided two alternative baselines of topological change introduced by increasing taxon sampling, and so the relative effect of these was compared to that of fossils by fitting separate linear models. However, as already explained, this approach can be problematic as the proportion of missing data differs between these types of terminals (Fig. S1 of SI File 3). To more directly account for this, a second analysis was performed using the amount of added information (i.e., the sum of the number of coded characters across added terminals, also log-transformed) as the continuous predictor in the linear model. Given that this approach fully accounts for differences in the amount of coded characters, only data for fossil and extant terminals was analyzed in such a way.

In each case, models with and without an interaction term were further compared using likelihood ratio tests (LRTs) with a significance level (α) of 0.1. In case of a significant interaction term, regions of significance were estimated using the Johnson-Neyman approach with package *jnt* (Middleton 2016); otherwise results presented are those of ANCOVAs. The Johnson-Neyman procedure finds the values of the continuous predictor for which there is a transition from significant to non-significant differences among the groups of the categorical predictor. In cases of non-significant differences between the different types of added terminals, a power analysis using the method of Borm et al. (2007) was performed (with a standard power level of 0.8).

Finally, we also relied on the construction of treespace (Hillis et al. 2005) to explore the type of topological change exerted by the addition of extant and fossil taxa. This approach is based on estimating all pairwise RF distances and decomposing them into a two-dimensional space using principal coordinate analysis (PCoA) (Jombart et al. 2017), providing a summary of the diversity of optimal topologies that are obtained as different types of terminals are incorporated into the inference procedure. Given the combinatorial nature of this method, we further subsampled at most 20 trees per analysis, and did not include topologies obtained after the addition of pseudoextinct terminals. Different treespaces were built for each combination of dataset, inference method and initial replicate. After obtaining the location of each tree in the first two PCoA axes, sets of trees from different phylogenetic analyses were collapsed to their centroid, and the relative position of centroids generated by the addition of fossil and extant terminals were compared using convex hulls and permutational (non-parametric) MANOVAs. Convex hulls provide a straightforward way in which to estimate the degree to which fossil and extant taxon addition results in the exploration of similar regions of treespace. Topological similarity was expressed as the area of overlap between the two convex hulls divided by the sum of the total area covered by the two, calculated using package *geometry* (Habel et al. 2019). Permutational MANOVA (Anderson 2001) is an analysis of variance using distance matrices, and employs a permutational test to evaluate significant sources of variation. The test was run for each treespace individually with package *vegan* (Oksanen et al. 2019), using Euclidean distances and evaluating significance with 10,000 replicates.

Given that the 25 initial replicates of each dataset do not share the same taxonomic composition, and therefore cannot be placed in a common treespace, a visual summary per dataset was obtained by using Procrustes superimposition. The different treespaces of each dataset were rotated so as to minimize the sum of squared distances between centroids obtained after adding the same number and type of terminals across replicates. In order to further facilitate interpretation of the plots, the different treespaces were then translated so that the average position of trees produced before taxon addition was at the origin.

### Estimating and Analyzing Taxon Influence

We also analyzed the topological effect induced by individual taxa using first-order taxon jackknifing. Unlike previous efforts however (Cobbett et al. 2007; Mariadassou et al. 2012; Denton and Goolsby 2018), we first standardized the size of the matrices from which taxa were jackknifed. We consider this approach to be superior for multiple reasons. First of all, it is evident from our results (as well as from previous analyses; e.g., Huelsenbeck 1991; Wiens 2003b, 2005; Prevosti and Chemisquy 2010) that the topological impact of a taxon will strongly depend on the size of the dataset to which it is added (see Results). If differences in size across datasets are not accounted for, these are bound to distort the impact of taxa when combining results from different datasets. Furthermore, standardizing matrix size has the added benefit that the impact of taxa can be measured against different taxonomic backgrounds. The resulting average impact is likely to be a better predictor of the effect a taxon will exert if incorporated into a new morphological matrix. Finally, it also allows for more intense replication and increased statistical power, while avoiding prohibitive computational times.

For each taxon, 30 matrices composed of the outgroup and 38 other randomly selected ingroup taxa were built. Unlike the approach described above, this subsampling scheme produces matrices composed of a mix of extinct and extant terminals. Phylogenetic inference was performed before and after the addition of the focal taxon to each of these matrices. A total of 27,930 MP and BI analyses were performed. Both phylogenetic inference and estimation of topological impact were performed as explained above. The impact of taxa under BI and MP was significantly correlated (Pearson’s r = 0.40, *p* < 10^−16^, Figs. S3 of SI File 3). Topological impact was then averaged across replicates to obtain a single measure per taxon. Given that we found a very strong structuring of topological impact by dataset (Figs. S4 and S5 of SI File 3), we subtracted from the estimated impact of each taxon the average impact for its dataset of provenance. This rescaling produces a metric that expresses deviations from an average impact and accounts for the difference in stability across datasets.

Generalized linear models were again employed to survey potential determinants of topological impact. We attempted to quantify all possible determinants previously hypothesized in the literature to affect a taxon’s topological impact. Some of these properties were estimated directly from the morphological matrices, while others were measured on BI posterior topologies (as they required branch lengths). From the matrices, we measured the proportion of missing/inapplicable data and two measures of morphological distance, which we refer to as ‘distinctiveness’ and ‘uniqueness’. These correspond to the minimum and average morphological distance to all other taxa, respectively, and were estimated using the maximum observable rescaled distance (Lloyd 2016). Among the tree-based metrics we quantified—for each focal taxa—the patristic distance to the root of the tree (‘primitiveness’), the length of its terminal branch (‘autapomorphies’), as well as three measures designed to capture different conditions that might lead to long-branch attraction scenarios. These included the change induced by the focal taxon on the mean and variance of branch lengths, as well as the variance of root-to-tip distances (‘clock-likeness’). Models also included the taxon type (fossil/extant) as an additional predictor. Further details on how these measures were quantified can be found in SI File 2. In order to explore the relevance of these predictors, stepwise variable selection in both directions was performed, once again comparing models using LRTs with α = 0.1.

Finally, we also explored the determinants of topological impact using random forest models. Random forests were chosen given their robustness to multicollinearity and deviations from normality, as well as their ability to accommodate non-linear and local relationships. Both the entire set of independent variables and the subset that can be estimated without running a phylogenetic analysis (i.e., all three matrix-based variables and the taxon’s type) were employed. These were used as predictors of both the topological impact score (regression forest), as well as a recoded binary variable grouping taxa into those with above and below average impact (classification forest). Models were fit using package *randomForest* (Liaw and Wiener 2002), growing 5,000 trees and estimating accuracy of prediction using out-of-bag errors. The number of predictors tried at each step was automatically determined by the algorithm.

R code to perform all of the procedures detailed above (other than phylogenetic inference) and perform the statistical analyses is available as SI File 5. It can be used to replicate all stages of the study alongside SI File 1 (original datasets) and SI File 6 (which contains all results).

## RESULTS

Our results show that the average degree of induced topological change strongly depends on the logarithm of the amount of information added to the analysis, measured as either the number of taxa or the number of coded cells incorporated to the matrix (Figs. 2, 3 and S6 of SI File 3; linear regressions, all R^2^ > 0.86, *P* < 8e-5). As the taxonomic sampling of phylogenetic analyses increases, it becomes progressively less likely that new data will further modify tree topology. The relative effect of adding new extant or fossil terminals proved highly dependent on the dataset, method of inference and number of added taxa (Fig. 2), although there was an overall stronger evidence for the addition of extant taxa generating higher topological impacts. Six of twelve analyses performed (i.e., combinations of datasets and inference procedures) showed a significantly higher impact of extant taxa across most/all of the range analyzed, while the opposite was only true for one analysis (echinoid dataset under BI; see also Fig. S6 of SI File 3). In stark contrast, fossil terminals had a significantly higher topological effect than pseudoextinct terminals, across all matrices and methods of inference (Fig. 3). This discordance reveals the strong relationship between topological impact and levels of missing data, and questions whether fossil and extant terminals are directly comparable.

**Figure 2:**
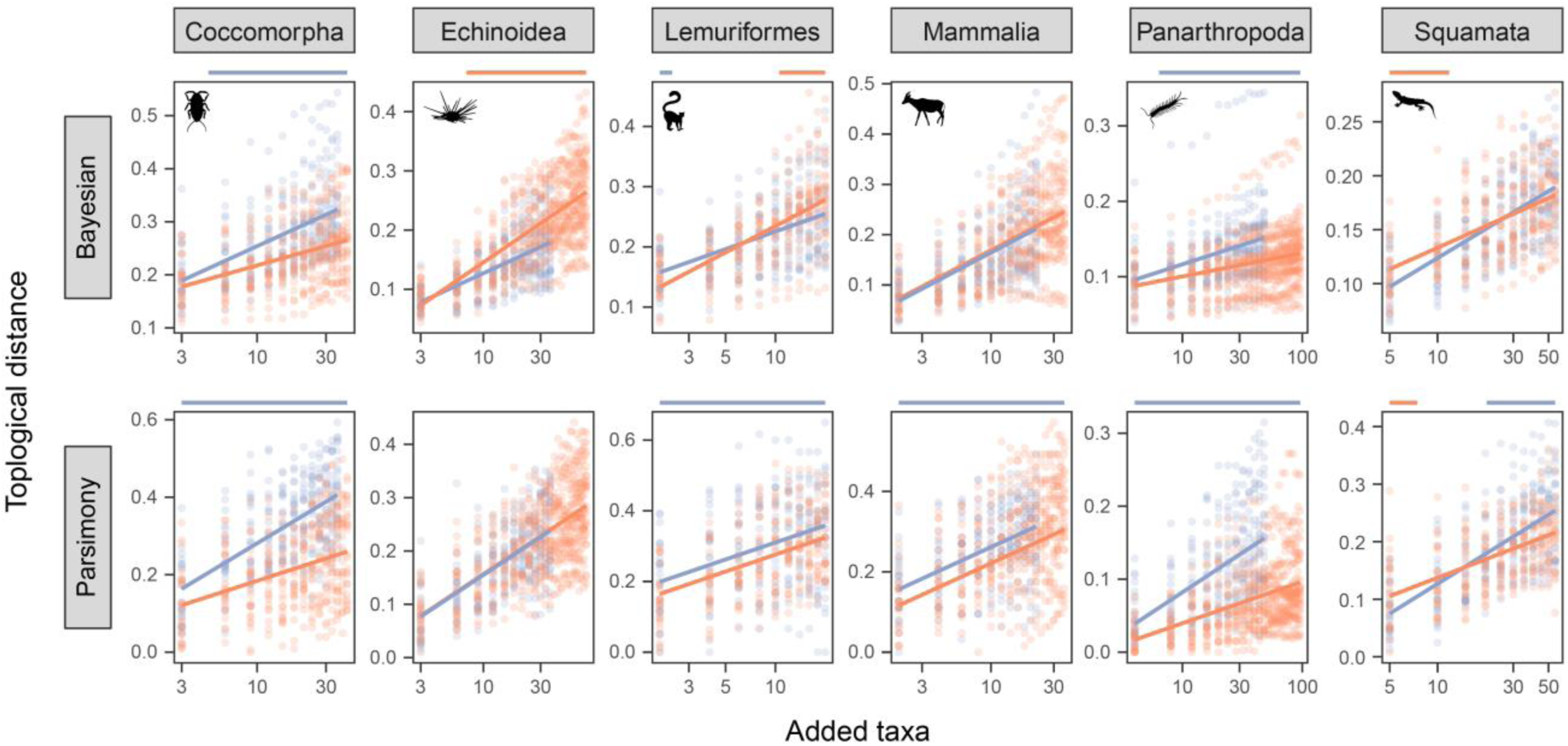
Comparison of the topological impact of fossil (orange) and extant (blue) terminals. Each dot is the topological distance (average RF distance to nearest neighbor) between trees before and after the addition of groups of either extant or fossil terminals, averaged across iterations. Regression lines are shown for the best-fit linear model. When the lines are parallel, the interaction term was not significant. Regions of significance are shown above each plot (the entire range is marked in the absence of a significant interaction). Silhouettes were taken from PhyloPic (http://phylopic.org/), and will be used throughout.

**Figure 3:**
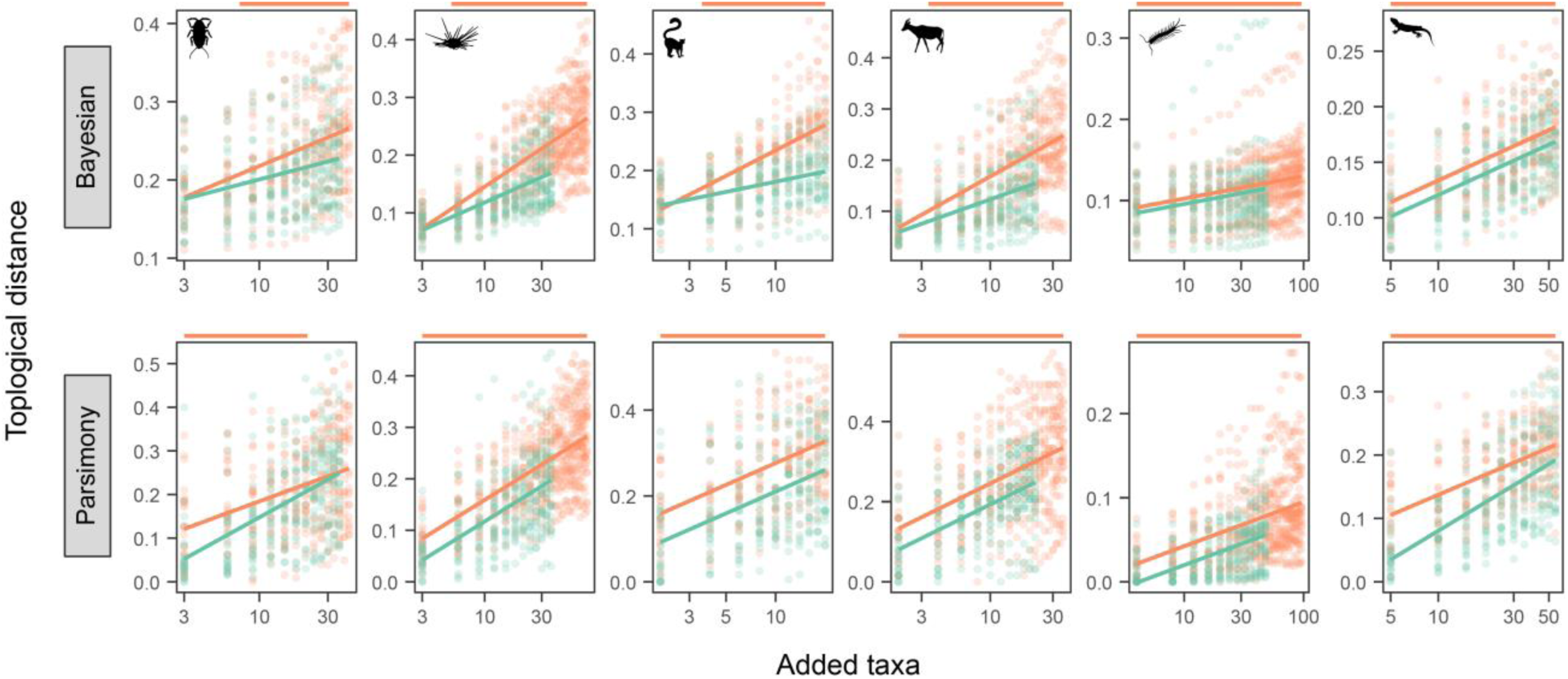
Comparison of the topological impact of fossil (orange) and pseudoextinct (green) terminals. Analysis of the data is performed as in Figure 2.

For this comparison to be meaningful, statistical models need to account for the fact that fossil and extant terminals do not contain equivalent amounts of information (see Table 1 and Fig. S1 of SI File 3). One possible way of achieving this is to compare the topological effect induced by the addition of paleontological and neontological data (rather than the number of terminals) to the inference procedure. We defined this variable as the sum of the number of characters that were neither missing nor inapplicable across the added terminals. From this perspective, differences in the behavior of these types of data are evident, with four matrices showing a stronger topological impact of paleontological data and two (Coccomorpha and Panarthropoda) a stronger effect of neontological data (Fig. 4). No systematic difference was found between the results obtained for MP and BI, which were very similar across datasets. The sole difference found was the behavior of the different types of data for the lemuriform dataset, which proved significant under BI but not under MP. This likely stems from a lack of statistical power, as this is the dataset with the smallest number of taxa, and therefore the fewest datapoints. In fact, a power analysis revealed significance should be attained with an increase of 16% in sample size. The lack of significance is therefore not given much weight, especially as the relative effects of paleontological and neontological data are the same for both inference procedures.

**Figure 4:**
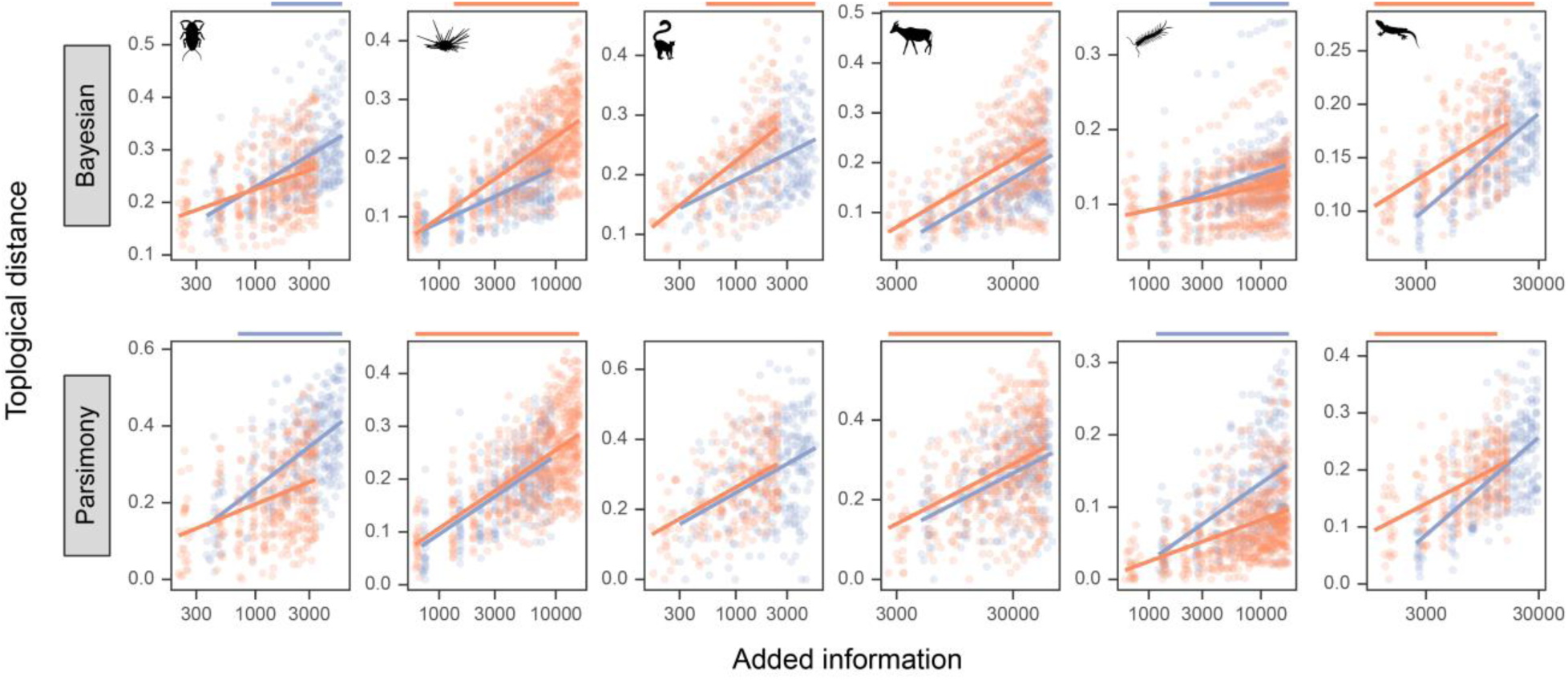
Comparison of the topological impact of paleontological (orange) and neontological (blue) data. Analysis of the data is performed as in Figure 2, although the continuous predictor employed is the logarithm of the amount of information added to the inference procedure (averaged across iterations).

We further compared the generated topologies through the use of treespaces, allowing us to explore the type of topological change induced by fossils and test if it systematically differed from that generated through the addition of further extant terminals (Fig. 5). As the trees obtained from the multiple replicates performed on each dataset do not share the same terminals (see Fig. 1 and Materials & Methods) and therefore cannot be represented in a common treespace, each panel of Figure 5 consists of the optimal juxtaposition of multiple distinct treespaces. The almost complete lack of overlap between the clouds of dots seen in Figure 5 is therefore even more pronounced if individual treespaces are scrutinized individually. Among these, convex hulls for topologies obtained after extant and fossil addition were completely disjunct 19.3% of the time, and otherwise showed an average overlap of only 9.8% of the total area covered. In fact, the relative positions occupied by these topologies were significantly different in 95% of the cases (permutational MANOVA, *p* < 0.05), with 13 of the 15 non-significant differences restricted to the MP analysis of the panarthropod dataset.

**Figure 5:**
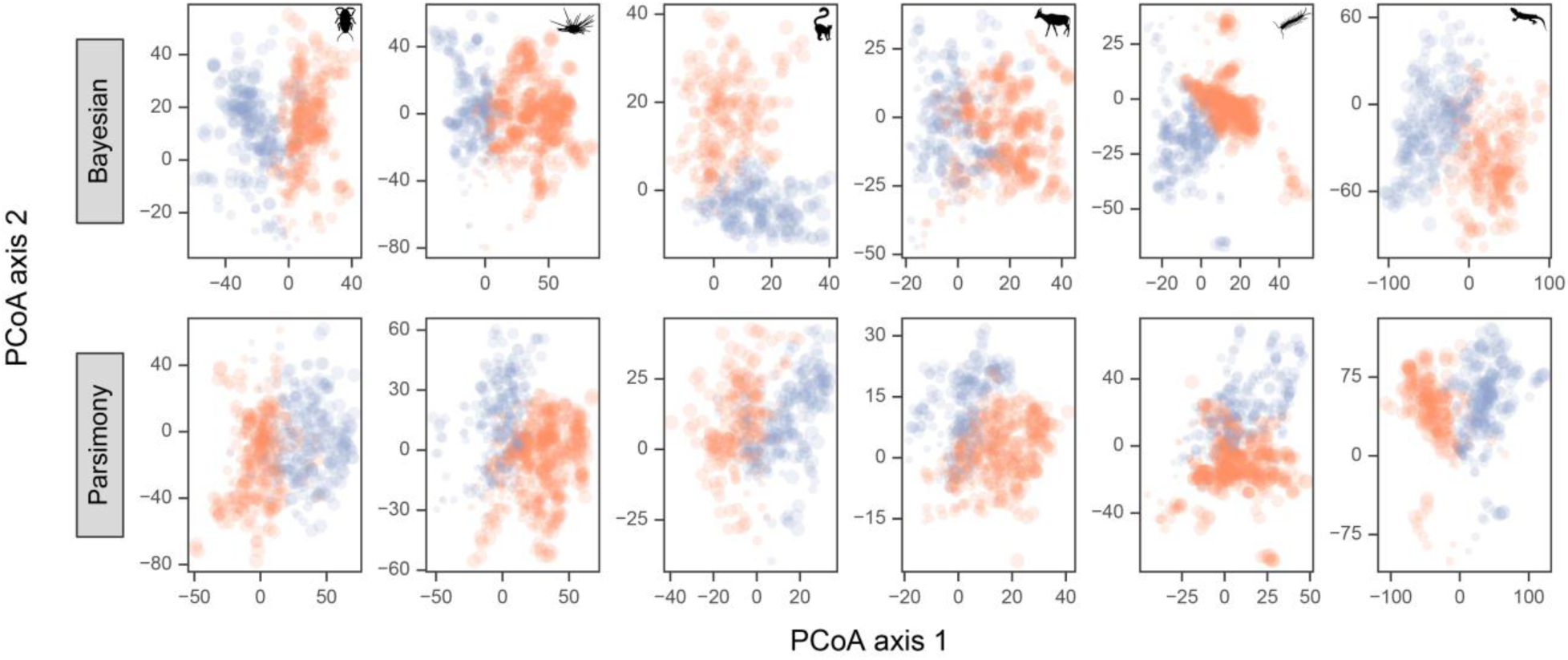
Topological changes induced by the addition of paleontological and neontological data to trees of extant terminals. A single treespace per replicate was generated, summarizing the position of trees obtained after incorporating increasing numbers of extant (blue) and fossil (orange) terminals. Treespaces for each dataset were then rotated to maximize superimposition, and translated so that the topologies product of the initial replicate (that obtained before any terminal is added, see Fig. 1, step 2) is at the origin. Dot size scales with the number of added terminals.

It is evident from this suite of analyses that fossil taxa have a unique behavior in phylogenetic inference (Figs. 2-5). However, this result raises the question of what determines the topological impact of a taxon. To address this question, we resorted to first-order taxon jackknifing experiments, extending previous efforts by assessing the topological effect of each terminal on multiple smaller matrices of the same size and composed of randomly sampled extant and fossil taxa (see Materials & Methods). In complete agreement with previous results (Cobbett et al. 2007), we found very small differences between fossil and extant terminals in their relative effects on topology. The type of terminal (fossil/extant) was not a significant predictor of topological impact for BI (*t*-test, *p* = 0.91), while fossils had a slightly smaller than expected impact under MP (*p* = 0.04). This result was mostly driven by the mammal dataset, the only individual matrix for which differences were significant (*p* = 0.001). However, this approach once again relies on comparing groups of terminals that systematically differ in their proportion of missing data (Table 1, Fig. S1 of SI File 3). After accounting for this difference, fossils were shown to impact topology significantly more than extant terminals above a threshold of 43-45% missing data (depending on the method of inference, Fig. 6a and Fig. S7 of SI File 3), a condition satisfied by over 78% of fossils in our dataset. Extant taxa are expected to have a significantly higher impact than fossils only in the very narrow condition of no missing data (although this requires extrapolation, and holds true only for MP, see Fig. S7 of SI File 3).

**Figure 6:**
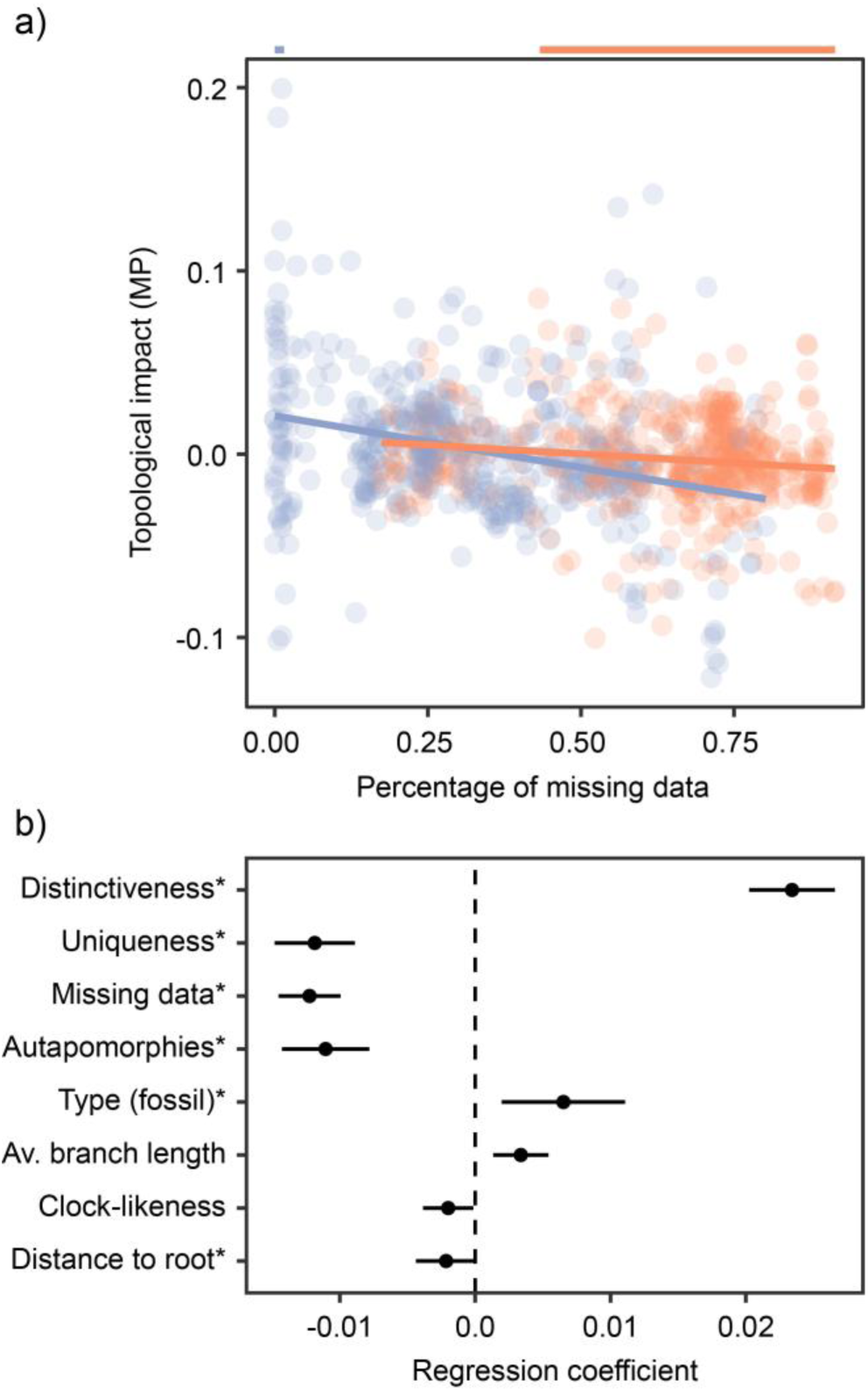
Exploring the determinants of topological impact under MP. **a)** Both missing data, type of taxa (fossil/extant) and their interaction plays a role in determining the topological effect of individual terminals. Across most of the region of missing data for which extant (blue) and fossil (orange) taxa overlap, fossils have a significantly higher topological impact (see also Fig. S7 of SI File 3). **b)** Effect size of the significant determinants of topological impact. Variables are ordered (top to bottom) according to the order in which they are incorporated into the stepwise linear model. Variables with an asterisk were also found to be significant determinants under BI (Fig. S8 of SI File 3). Effect sizes are standardized to reflect expected changes generated by a difference of one standard deviation.

We quantified several other matrix and tree-based properties of terminals and explored their usefulness as predictors of topological impact using generalized linear models. The obtained best-fit models under BI and MP were extremely similar, yet explained widely different proportions of total variance (adjusted R^2^ of 0.06 and 0.31, respectively). For this reason we focus here only on the results for MP. All of the results obtained under BI are placed in SI File 3, and a discussion of the differences between methods can be found in SI File 2.

Our analysis revealed the presence of atypical morphologies as a major driver in a taxon’s topological impact (Figs. 6b and S8 of SI File 3). Organisms displaying character state combinations distinct from those of all other taxa strongly modified topology, while completely unique morphologies had a significantly lower impact. The amount of missing data and the degree of evolutionary change also had negative effects, with taxa displaying ‘primitive’ morphologies (i.e., those inferred to have undergone the least amount of change since the last common ancestor of the ingroup) and possessing fewer autapomorphies having stronger effects on topology. Furthermore, terminals that increased the average branch length of trees and decreased the clock-likeness of morphological evolution also showed a higher impact. It is possible that these taxa are generating conditions conducive to long-branch attractions. Finally, even after accounting for the aforementioned variables, fossils still induced stronger topological changes than extant taxa, suggesting that the peculiarities of paleontological data are still not completely captured by our model.

The topological impact of terminals has been claimed to be unpredictable (Cobbett et al. 2007), yet to our knowledge no attempt at directly predicting it has been performed. Given how random forest models using the same set of predictors were able to explain an even higher proportion of variance than linear models (44% in the case of MP, see Table S1 of SI File 7), we tested the accuracy of random forests to predict whether a taxon had a higher or lower-than-average topological impact. Prediction was carried out using only the set of four variables that are tree-independent and can therefore be estimated before running a phylogenetic analysis (distinctiveness, uniqueness, proportion of missing data and type of taxon). Even using this reduced set of variables, the model attained a classification accuracy of 72% (Table S2 of SI File 7), indicating that the topological impact of taxa in morphological phylogenetic analysis has a strongly predictable component.

## DISCUSSION

For decades, systematists have disagreed on the relevance that the fossil record has for inferring phylogenetic relationships. By focusing on just a few of the many ways in which paleontological and neontological data differ, fossils were portrayed to be either vital or irrelevant for phylogenetic inference, as well as everything in between. However, much of this discussion was based on either first principles or individual case studies. The only attempt to systematically assess the relevance of fossils suggested that differences in the behavior of extant and extinct taxa might have been exaggerated, and that fossils do not differ systematically from extant terminals, at least not in terms of their average topological effect.

Our results are able to reconcile these seemingly incompatible claims about the nature of the fossil record and its relevance for systematics. All of our analyses reinforce the idea that the information preserved in the morphology of extinct organisms is unlike data that can be obtained from the study of living species, and that its inclusion in phylogenetic inference has strong consequences. However, this is evident only after the fragmentary nature of fossils is accounted for. Missing data does in fact affect topological impact (Fig. 6a), something that had already been demonstrated through simulations (Huelsenbeck 1991; Wiens 2003b, 2005), although not always confirmed using empirical datasets (Denton and Goolsby 2018). While previous studies had directly compared fossil and extant taxa, a better understanding of their behavior required accounting for the different amount of information they contain. When this is done, paleontological data tends to have stronger topological effects than neontological data (Figs. 2-4, 6, S6-S8). Furthermore, adding fossils induces topological changes that are never obtained through better taxonomic sampling of extant taxa (Fig. 5), a phenomenon that likely underlies their ability to resolve conflicts between morphological and molecular phylogenies (Legg et al. 2013; Beck and Baillie 2018; Simões et al. 2018; Asher et al. 2019).

Why do fossils show such a remarkable effect on phylogenetic analysis? Our results show morphological distinctiveness to be the main predictor of topological impact (Figure 6b). The fossil record contains a wealth of examples of extinct organisms with morphologies wholly unlike those we see in the modern biosphere, and whose appearance we would not be able to predict even with the closest scrutiny of the morphology of extant taxa. Examples of such organisms are found throughout geological time, from the ‘weird wonders’ of the Cambrian to the diversity of extinct archosaurs that populate the stem lineage of birds. The unique combination of characters in such organisms has long been recognized to modify character polarity, reveal hidden homoplasies (or cryptic homologies) and break long branches by revealing the sequence of character acquisition (Gauthier et al. 1988; Donoghue et al. 1989; Smith 1998; Edgecombe 2010). However, taxa with completely unique morphologies exhibit decreased topological impact, likely attaching to a topology without modifying character optimization. In such cases of extreme morphological modifications, such as those separating hexapods and crustaceans (Legg et al. 2013), or cetaceans from their ungulate relatives (Spaulding et al. 2009), fossil stem groups with intermediate morphologies are required to correctly articulate these clades. Finally, the fossil record provides our only access to morphologies that lie close to the origin of ancient lineages and are less burdened by character state changes imposed by subsequent evolutionary history, properties that further increase the topological impact of individual taxa.

Fossils not only play a critical role when inferring evolutionary processes from phylogenetic trees (Slater et al. 2012; Goswami et al. 2016), but are clearly also influential in the construction of the trees themselves. Beyond recognizing the unique role of paleontological data in phylogenetic inference, our analyses prove that topological impact has strong determinants. Consideration of these properties can provide new ways for optimizing the phylogenetic design of total-evidence analyses, aiding in the generation of phylogenies that better incorporate the vast records of evolutionary history preserved in the morphology of extant and extinct lineages. Given that genomic data is yet to resolve a stable phylogeny for all branches in the tree of life, phylogenetic analyses of morphology incorporating fossil taxa remain an important and independent framework for unraveling evolutionary history.

## SUPPLEMENTARY MATERIAL

Data available from the Dryad Digital Repository: http://dx.doi.org/10.5061/dryad.[NNNN].

## ACKNOWLEDGEMENTS

This manuscript was enriched by discussions with Jacques A. Gauthier, Michael J. Landis, Michael J. Donoghue and Derek E. G. Briggs. We would also like to thank Kaylea Nelson for computational assistance.

## AUTHOR CONTRIBUTIONS

NMK designed the study and wrote all code. NMK and LAP gathered the original datasets and ran the analyses. NMK analyzed the data and wrote the manuscript with contributions from LAP.

