## Supplementary material for "Death is on Our Side: Paleontological Data Drastically Modify Phylogenetic Hypotheses": SI File 2

**SI File 2: Supplementary Methods**

**Sections:**

I. Dataset selection and processing
II. Subsampling procedure
III. Determinants of taxon influence
IV. Comparison of results between parsimony and Bayesian inference
I. Dataset selection and processing

Datasets were obtained from the literature, downloaded from online repositories, or obtained directly from the authors. For some of these, taxonomic sampling was markedly uneven across clades, and so the ingroup (IG) was redefined as the clade with the most extensive taxon sampling. The composition of the IG follows that of the source of the dataset. In each case, a single outgroup (OG) was included to root all trees, chosen for being either the only representative of the sister lineage to the IG, or the OG terminal with the least amount of missing data if multiple terminals were sampled from the IG’s sister clade. Datasets for Coccomorpha (Vea and Grimaldi 2015), Panarthropoda (Legg et al. 2013) and Squamata (built upon the datasets of Gauthier et al. 2012 and Mongiardino Koch and Gauthier 2018, although largely unpublished) were not modified in any other way. The remaining matrices were modified as follows:

- Echinoidea (Kroh and Smith 2010): The version of this dataset employed is not the original but rather that of Hopkins and Smith (2015), which has five fewer IG taxa. These were pruned due to their unstable phylogenetic position (as also noted by the original authors).
- Lemuriformes (Herrera and Dávalos 2016): This dataset had markedly uneven sampling between strepsirhine and haplorhine primates (83% vs. 3% of recognized extant species, as reported by the authors). We therefore redefined the IG as crown-group strepsirhines (i.e., Lemuriformes following the nomenclature of Szalay and Delson 2013), pruning all haplorhines and stem primates (according to the topology supported by this dataset, see Figure 3 of the original publication).
- Mammalia (O’Leary et al. 2013): The dataset deposited in MorphoBank (<http://dx.doi.org/10.7934/P773>) contains 12 characters with more than 10 states, which is the maximum number of states for morphological characters accepted by MrBayes. Therefore, states above this limit were recoded as missing data, as was also done by the authors (see Supplementary Materials of the original publication). For consistency, both maximum parsimony (MP) and Bayesian inference (BI) analyses were performed with this modified matrix rather than retaining the original scores for parsimony analyses.

Edited datasets are available as SI File 1, and their final sizes can be found in Table 1. Ordered characters (present only in the echinoid and squamate datasets) and polymorphisms were analyzed as such, and no distinction was made between missing and inapplicable data.

II. Subsampling procedure

Datasets were imported into R (R Core Team 2019) using function ReadMorphNexus from package Claddis v. 0.3 (Lloyd 2016). These were then subsampled by selecting a number ($n$) of extant taxa at random (see Figure 1, step 1). The value of $n$ was different between datasets, and was chosen depending on the absolute number of fossil ($f$) and extant ($e$) terminals available. Taking into account these numbers guaranteed that topological changes could be reliably assessed across datasets (i.e., initial topologies had multiple internal nodes while leaving large and comparable numbers of fossil and extant terminals unsampled). If $e>2*f$, then $n=\left( e+f \right)-2*f=2*e-(e+f)$. This condition leaves the same number (*f*) of both fossil and extant taxa unsampled from the initial dataset. Otherwise, if $e<2*f$, then $n=e/2$. Under this condition the number of fossils exceeds that of unsampled extant taxa. To these initial replicates, different types of terminals were added in a stepwise manner in groups of size $m$ until unsampled terminals were exhausted. Only in the case of fossil addition to the panarthropod dataset was the procedure stopped before exhausting terminals (at a final matrix size three times that of the initial replicate). The value of $m$ was chosen to guarantee at least 10 steps of taxon addition, and varied between 2 and 5 depending on the dataset. The actual numbers used for each dataset can be found in SI File S2.

III. Determinants of taxon influence

The determinants of a taxon’s topological influence included some properties that were estimated directly from the morphological matrices, as well as others that were measured on Bayesian inference (BI) posterior topologies (as they rely on the estimation of branch lengths). The matrix-based properties included the proportion of missing/inapplicable data and two measures of morphological distance, which we refer to as ‘distinctiveness’ and ‘uniqueness’. These correspond to the minimum and average morphological distance to all other taxa, respectively, and were estimated using the maximum observable rescaled distance (MORD; Lloyd 2016). This distance is similar to Gower’s coefficient (Gower 1971), but is standardized relative to the maximum realizable distance given the structure of the dataset, instead of using the number of variables for which pairs of observations can be compared. When datasets have ordered characters (as in some of the datasets employed) these two distances can differ. Several benefits of using MORD over Gower’s coefficient when dealing with morphological matrices have been described (Lloyd 2016; Lehmann et al. 2019). Furthermore, unlike Gower’s coefficient, MORD produces values that are scaled between 0 and 1, thus allowing for more straightforward combination of data from different datasets. Our two variables capture different aspects of morphological variability. For example, highly distinct taxa (i.e., those having uncommon character-state combination) are bound to be located on relatively unexplored regions of morphospace, but they might not be highly unique if there is a second taxon with a similar morphology (i.e., the distance to the nearest neighbor can still be relatively small). These two aspects of the morphological variability present in the dataset are likely to have different consequences for a taxon’s topological impact. Distances were not used for ordination and hence were left untransformed (Lloyd 2016).

All tree-based metrics were quantified in the set of 100 randomly subsampled trees obtained from each BI analysis and averaged. For each focal taxon (i.e. the one being jackknifed) we measured:

- The patristic distance between the taxon and the root of the tree. We used this metric as a proxy for the ‘primitiveness’ of the terminal, as it captures the amount of expected morphological change its lineage has undergone since the last common ancestor of the IG. Calculation relied on function distRoot from package *adephylo* (Jombart and Dray 2010).
- The length of the terminal branch leading up to the taxon. We used this as a proxy for the number of autapomorphies the terminal has accrued since the last cladogenetic event.
- The change in the mean and variance of the distribution of branch lengths before and after the addition of the focal taxon.
- The change in the variance of root-to-tip distances generated by the addition of the focal taxon. Root-to-tip distances are a commonly employed proxy for the degree to which evolution has operated in a clock-like manner (e.g., Smith et al. 2018).

The first two of these metrics (‘primitiveness’ and ‘autapomorphies’) were measured directly on the set of trees produced by inference on matrices containing the focal taxon, and were divided by total tree length (i.e., the sum of all branches) for standardization. The remainder represents the difference in the parameter estimated in the sets of trees obtained before and after taxon addition. Two of the metrics (‘primitiveness’ and ‘clock-likeness’) required rooting trees using the OG, while all others were estimated with unrooted topologies. While the interpretation of the two first two metrics is rather straightforward, the last three (changes in ‘clock-likeness’, mean and variance of branch lengths) were designed to capture different conditions that might render analyses more or less susceptible to long-branch attraction artifacts. We caution however that some of these variables might be affected by the priors used in the Bayesian analysis, as well as the model used to correct for ascertainment bias in the datasets, and efforts should be made to improve upon these proxies.

IV. Comparison of taxon influence between parsimony and Bayesian inference

While method choice did not impact any of the results comparing the magnitude and type of topological change exerted by adding fossil and extant terminals to matrices composed only of extant taxa (Figs. 2-4, S6-8 of SI File 3), it did substantially affect our ability to detect determinants of taxon influence.

Firstly, the average topological impact of taxa was much more strongly structured by dataset for the BI analyses than it was for maximum parsimony (MP). While the dataset of provenance (used as categorical predictor in an ANOVA) explained a large portion of variance in topological impact under MP (R^2^ = 0.621, Fig. S4 of SI File 3), it explained almost all of the variance under BI (R^2^ = 0.987, Fig. S5 of SI File 3). Datasets are in fact expected to differ in their level of stability (i.e., how susceptible the results are to changes/increases in taxon sampling), something that might be related to both the idiosyncrasies of the clade’s evolutionary history and different aspects of matrix construction (e.g., coding strategies, taxon sampling, dataset size, etc.). Although too few datasets are used here to explore these relationships, it is worth noting that the least stable dataset (Coccomorpha) is also the one with the smallest number of characters, while the most stable (Mammalia) is the also the largest (see Table 1 and Fig. S5 of SI File 3). Nevertheless, the degree to which the dataset of provenance explains results under BI might be indicative that a large proportion of the observed variability stems from differences in chain convergence between datasets, and is therefore unrelated to the true taxon influence.

Despite this, results from the statistical analyses performed after this dataset effect is subtracted from MP and BI analyses are remarkably similar. Fossils are found to impact phylogenetic inference more than extant terminals above a very similar threshold of missing data (43.3% for MP and 45.2% for BI, see Figs. 6a and S7 of SI File 3). The determinants deemed to be significant for both inference methods are very similar (Figs. 6b and Fig. S8 of SI File 3), and they affect taxon influence in the same way. Furthermore, topological impact estimated under BI and MP are significantly correlated (*P* < 10^-16^, Fig. S2 of SI File 3), even though the correlation coefficient is not particularly high (Pearson’s r = 0.402). It should also be noted that the total amount of variance explained by these significant regressors is dramatically different between inference methods. While 30.5% of variance in taxon influence is explained by the best-fit linear model obtained for MP, only 5.8% is explained by the model obtained for BI. The same is true for random forest regression models (44.2% vs. 7.9%), although once again the ranking of determinants by relative importance were very similar between MP and BI (see Table S1 of SI File 7). This difference in explained variance might once again reflect convergence issues, or might be a consequence of the broader distribution of optimal topologies that BI outputs relative to MP (O’Reilly et al. 2016, Schrago et al. 2018, Smith 2019). In this last scenario, subsampling of topologies according to their posterior probabilities (instead of at random) might improve the accuracy with which taxon influence is estimated (Denton and Goolsby 2018), facilitating the study of its determinants.

Given this difference in explained variance, a random forest classification model was fit only to the results of topological impact under MP. This model provides an estimate of the accuracy with which taxa can be classified into those having either higher or lower-than-average impact, using only predictors that can be estimated directly from the matrix (i.e., missing data, distinctiveness, uniqueness and type). The results show a relatively high degree of classification accuracy (71.75%), with percentage of missing data and both morphological distance variables having similarly high levels of relative importance (Table S2 of SI File 7). This result shows that the topological impact of taxa in phylogenetic analysis under MP seems to be (to some extent) predictable. Although the aforementioned similarities in the results obtained for inference under MP and BI indicate that topological impact under both methods seems to be affected by the same underlying factors, confirming this and extending these results into predictive frameworks will require improving the way impact is estimated under BI. Ensuring convergence of chains, using a larger number of trees from the posterior distributions for calculations, and sampling trees according to their posterior probability are some of the ways in which this might be achieved.
