## Supplementary material for "Death is on Our Side: Paleontological Data Drastically Modify Phylogenetic Hypotheses": SI File 3

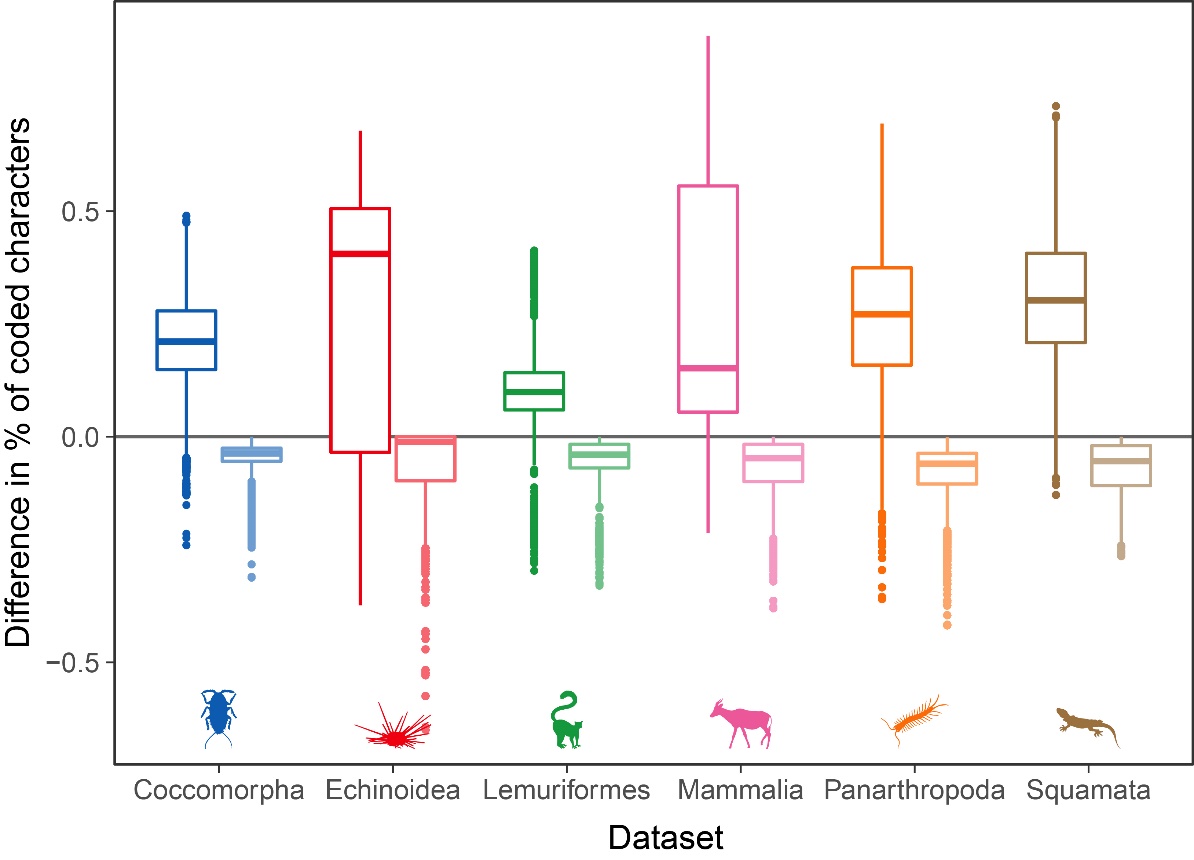

**Figure S1:** Differences in the percentage of coded characters between fossil and extant terminals (left, dark colors) and fossil and pseudoextint terminals (right, light colors). Values were obtained by randomly pairing terminals of each type and comparing the percentages of coded characters. The procedure was repeated 10,000 times for each pairing type and dataset. Given the differences in the proportion of missing data between these types of terminals, the direct comparison of the topological impact of extant and fossil addition (Figs. 2, S6) is likely biased towards favoring extant terminals. Similarly, the comparison of fossils and pseudoextinct terminals (Figs. 3, S6) is likely biased towards favoring fossil terminals, although the strength of the bias introduced by missing data is expected to be much lower in this case (2.3-4.6 times lower on average, depending on the dataset).

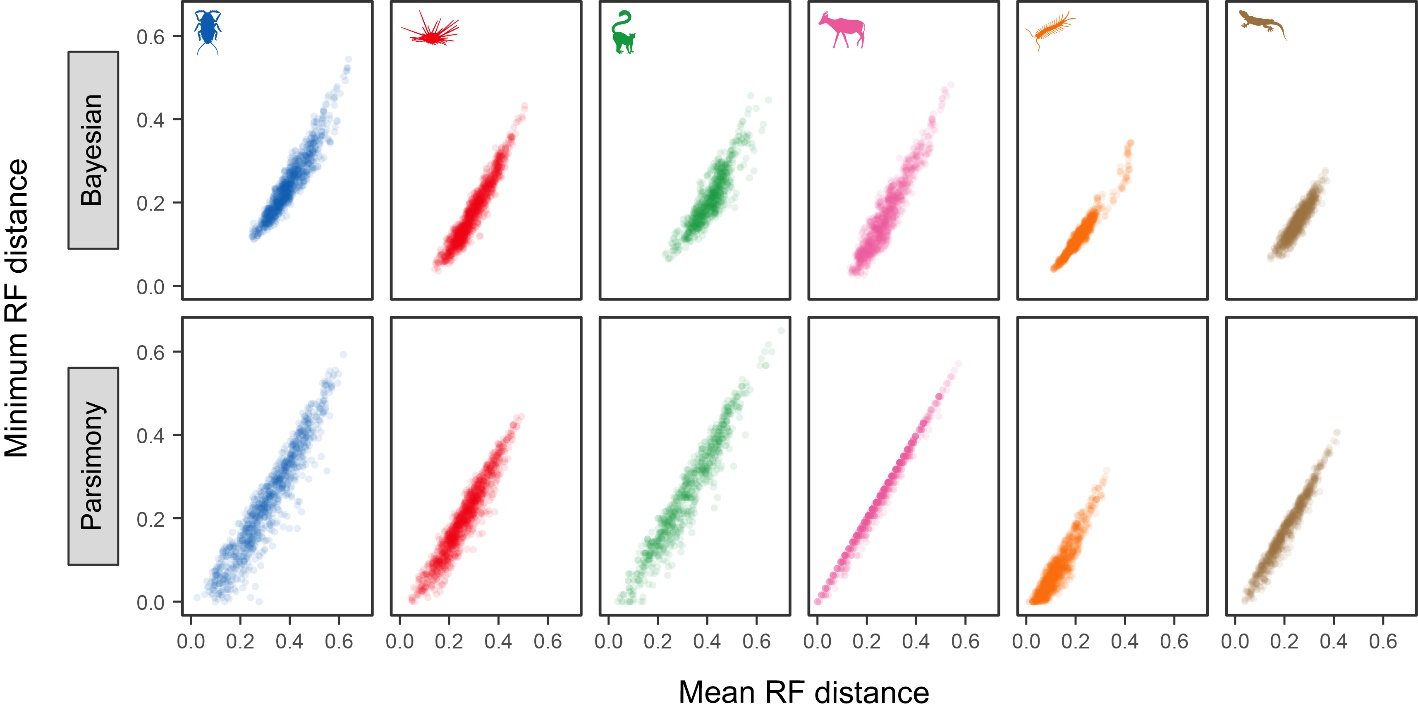

**Figure S2:** Alternative ways of calculating topological distances between sets of trees (average of the mean and minimum RF distances of each tree to all the trees in the other set) are very highly correlated. Choosing between them is not expected to modify downstream analysis, and therefore the minimum RF distance was employed.

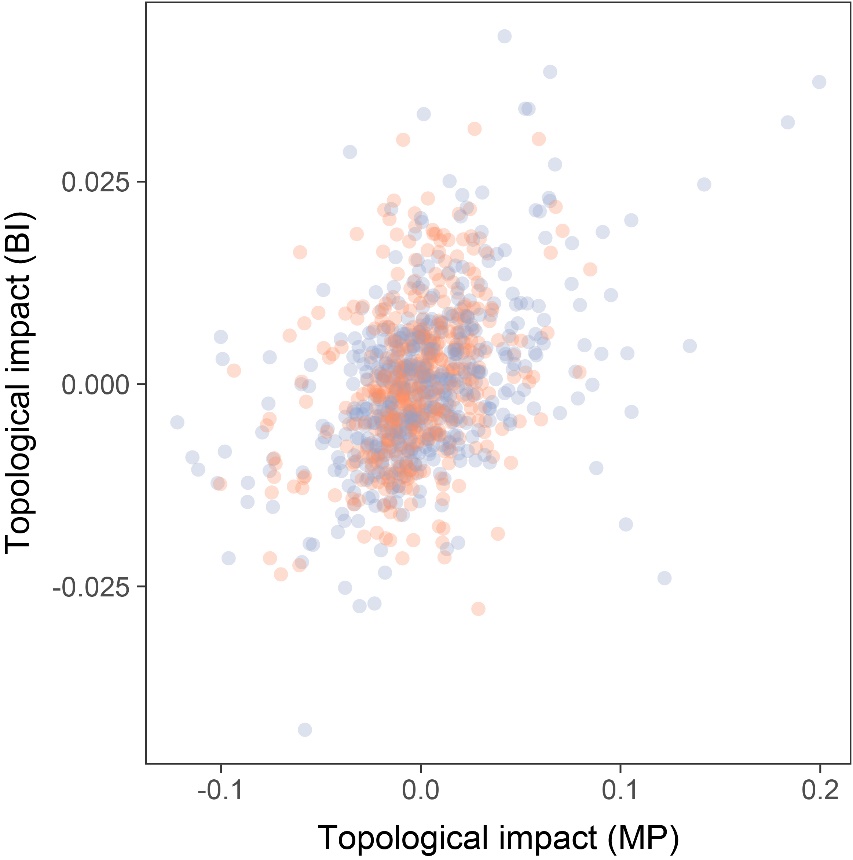

**Figure S3:** Comparison of the estimated topological impact of taxa under Bayesian inference and maximum parsimony. Orange dots correspond to fossils taxa, those in blue to extant taxa. Correlation is highly significant (*P* < 10^-16^) but not very strong (Pearson’s r = 0.402).

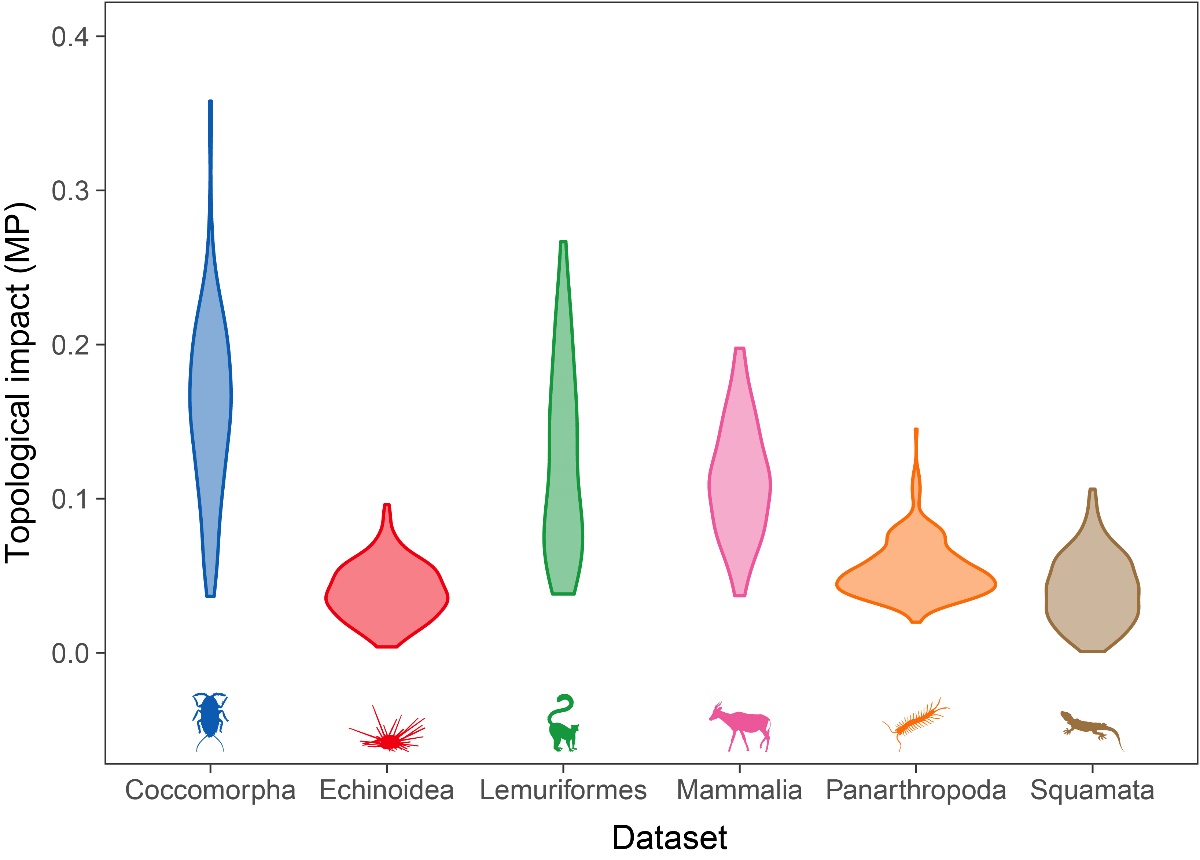

**Figure S4:** Distribution of estimated topological impact for terminals of different datasets under maximum parsimony (MP).

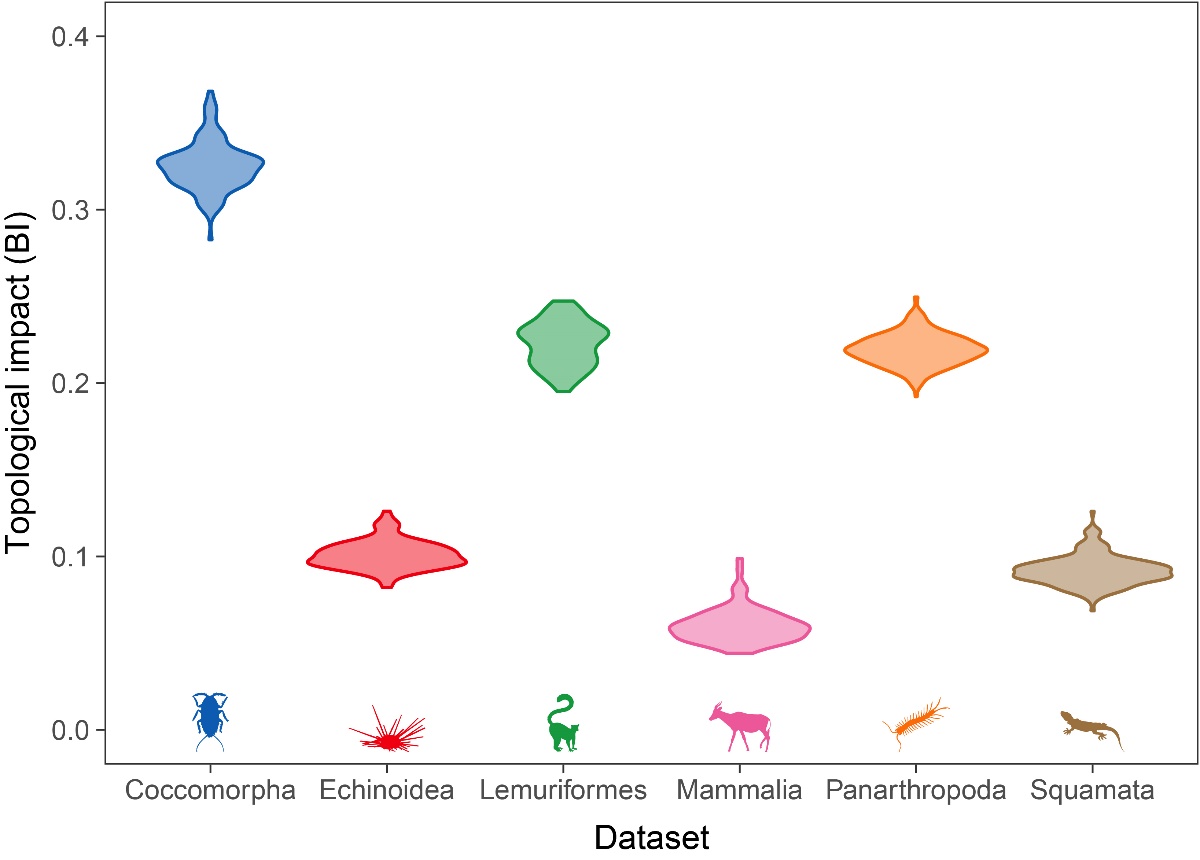

**Figure S5:** Distribution of estimated topological impact for terminals of different datasets under Bayesian inference (BI).

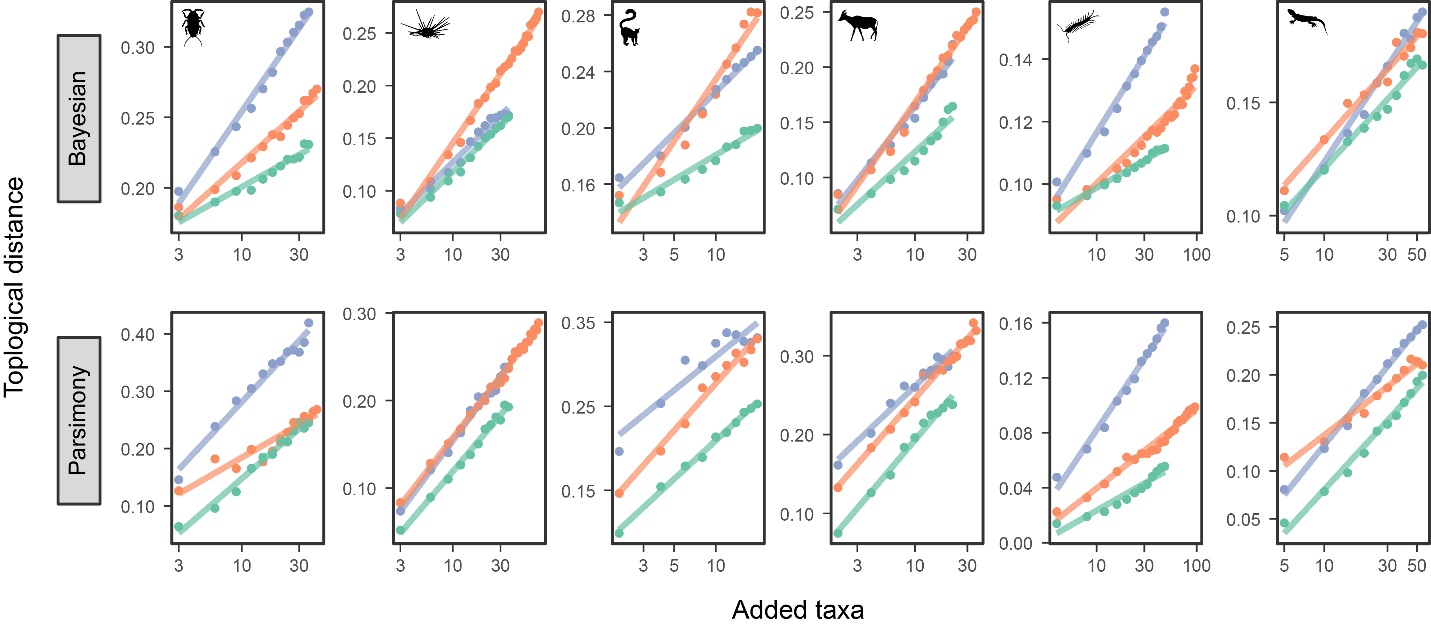

**Figure S6:** Comparison of the average topological impact generated by adding fossil (orange), extant (blue) and pseudoextinct (green) taxa to phylogenetic hypotheses built exclusively from extant terminals. This represents an overall summary of the data showed in Figures 2 and 3. Given the difficulty of visualizing all three treatments in a single plot, the data was further averaged to a single value for each combination of number and type of terminal added. Regression lines were estimated for each type of terminal independently.

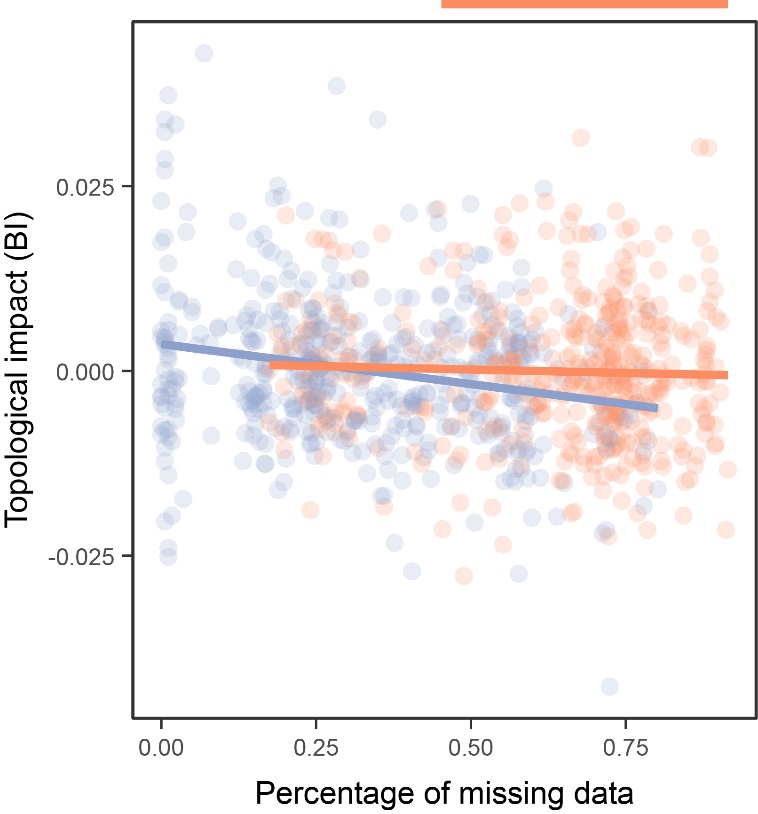

**Figure S7:** The topological impact under Bayesian inference (BI) of fossil (orange) and extant (blue) taxa, and its relationship with the percentage of missing data. Regions of significance are shown above the plot. Results are very similar to those shown in Figure 6a for maximum parsimony. However, note that there is no region of percentage of missing data for which extant taxa show a significantly higher impact.

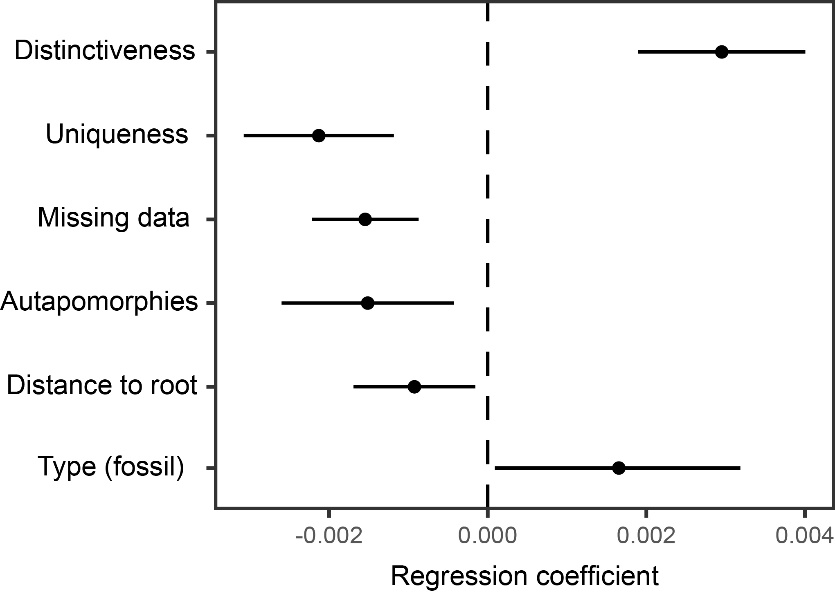

**Figure S8:** Effect size of the significant determinants of topological impact under Bayesian inference. Variables are ordered (top to bottom) according to the order in which they are incorporated into the stepwise linear model. Effect sizes are standardized to reflect expected changes generated by a difference of one standard deviation. Results are very similar to those shown in Figure 6B for maximum parsimony, although fewer variables are recovered as significant under Bayesian inference.
