## Supplementary material for "Death is on Our Side: Paleontological Data Drastically Modify Phylogenetic Hypotheses": SI File 7

**Table S1:** Relative importance of predictors in the random forest regression models built to explain differences in topological impact across taxa under MP and BI. Importance is measured using percentage increase in mean squared errors (%IncMSE). This is calculated as the difference in MSE on the out-of-bag portion of the data before and after randomly permuting each predictor variable. The difference between the two is averaged over all trees and normalized by the standard deviation of the differences (Liaw and Wiener 2002).

| **Predictor** | **%IncMSE for MP** | **Predictor** | **%IncMSE for BI** |
| --- | --- | --- | --- |
| Distinctiveness | 119.70 | Distinctiveness | 73.04 |
| Missing data | 108.83 | Missing data | 68.96 |
| Uniqueness | 94.93 | Clock-likeness | 68.63 |
| Primitiveness | 70.25 | Uniqueness | 61.63 |
| Clock-likeness | 68.38 | Av. branch length | 55.65 |
| Autapomorphies | 65.66 | Primitiveness | 55.50 |
| Av. branch length | 61.90 | Autapomorphies | 52.92 |
| Var. branch length | 43.26 | Var. branch length | 44.74 |
| Type | 24.88 | Type | 17.98 |

**Table S2:** Relative importance of predictors in the random forest classification model built to explain differences in topological impact across taxa under MP. Importance is measured as explained in the caption of Table 1, although using the difference in accuracy (classification error rate) before and after randomly permuting predictors (Liaw and Wiener 2002).

| **Predictor** | **Mean decrease in accuracy** |
| --- | --- |
| Missing data | 167.94 |
| Distinctiveness | 159.51 |
| Uniqueness | 149.57 |
| Type | 71.13 |
